# Early transcription factor activation distinguishes symbiotic from non-symbiotic bacteria during microbiome processing in a sponge

**DOI:** 10.1101/2025.05.27.656460

**Authors:** Bin Yang, Benedict Yuen, Huifang Yuan, Bernard M Degnan, Sandie M Degnan

## Abstract

Animals that filter-feed on environmental microbes must rapidly discriminate among captured bacteria to maintain beneficial associations while avoiding inappropriate immune activation. In innate immunity, this discrimination is executed through transcription factors (TFs), whose activation and nuclear translocation initiate effector gene expression and shape the nature of the host response. In sponges, bacteria are first physically captured by choanocytes, but the timing and cellular context in which TF-mediated immune discrimination becomes evident remains unclear. Here, we investigate the earliest detectable regulatory responses associated with discrimination between symbiotic and non-symbiotic bacteria in the marine sponge *Amphimedon queenslandica*. Using a feeding-based design to model post-metamorphic microbiome restructuring, we exposed juvenile sponges that already harbour vertically inherited symbionts to native (symbiont) or foreign (non-symbiont) bacterial communities and assessed early cellular processing and transcriptional responses to bacterial uptake. Symbiotic bacteria were rapidly transported across the epithelium and induced a strong, transient activation of conserved innate immune TFs, including NF-κB, IRF, and STAT, together with associated signalling pathways. IRF and NF-κB translocated to the nuclei of amoebocytes that had engulfed symbionts, indicating that discrimination becomes evident shortly after uptake and precedes downstream effector responses. In contrast, foreign bacteria were internalized more slowly, failed to induce coordinated immune TF activation or nuclear translocation, and instead elicited a xenobiotic-dominated transcriptional program. Together, these findings identify TF activation as an early regulatory checkpoint in sponge–microbe interactions and reveal key mechanisms that underpin the initial stages of symbiont discrimination.

## INTRODUCTION

The origin and evolution of animals occurred in the continual presence of bacteria, which long pre-dated metazoans in marine environments. Throughout their evolutionary history, animals have thus interacted continuously with bacteria in relationships ranging from antagonism to mutualism. These interactions likely imposed early selective pressure for mechanisms that distinguish beneficial from harmful microbes, in turn shaping the evolution of innate immunity [1–11]. Consistent with this, all animals share many components of innate immunity, including transcription factors (TFs), pattern recognition receptors (PRRs), signalling pathways, and core effector responses [3, 12–16]. In general, microbial encounters are translated into host responses via TF-mediated gene regulatory networks, with TF activation and nuclear translocation providing the critical regulatory step that initiates immune effector gene expression [11, 17–20]. Despite this deep conservation, how innate immune systems contribute to early discrimination among different types of bacterial associations remains poorly understood outside vertebrate systems [21–23].

Sponges provide a powerful system to address this gap. Their internal epithelial boundary comprises choanocyte chambers – flagellated epithelial cells that drive water flow and capture bacteria – and endopinacocytes that line internal canals [24, 25]. Beneath this epithelium lies a collagenous mesohyl containing phagocytic amoebocytes closely apposed to the basal surface of choanocytes, archaeocytes that are large pluripotent and multifunctional amoebocytes, and other secretory and biomineralizing cell types that interact with both food bacteria and symbionts [24–27] . As sessile suspension feeders, sponges continuously filter large volumes of seawater through choanocyte chambers, capturing diverse environmental bacteria [24, 25]. At the same time, many sponges maintain stable and often species-specific bacterial symbioses, with extracellular symbionts residing in the mesohyl, often near choanocyte chambers (Fig. 1A-C) [10, 28]. This ecology creates a persistent requirement to discriminate symbionts from the diverse array of transient, nutritional, or potentially harmful microbes encountered during feeding [29]. However, although choanocytes are the first cells to physically contact environmental bacteria, the molecular mechanisms operating at this interface remain poorly characterised, and it is unclear when transcriptional immune commitment first becomes evident during microbiome processing.

**Fig. 1.**
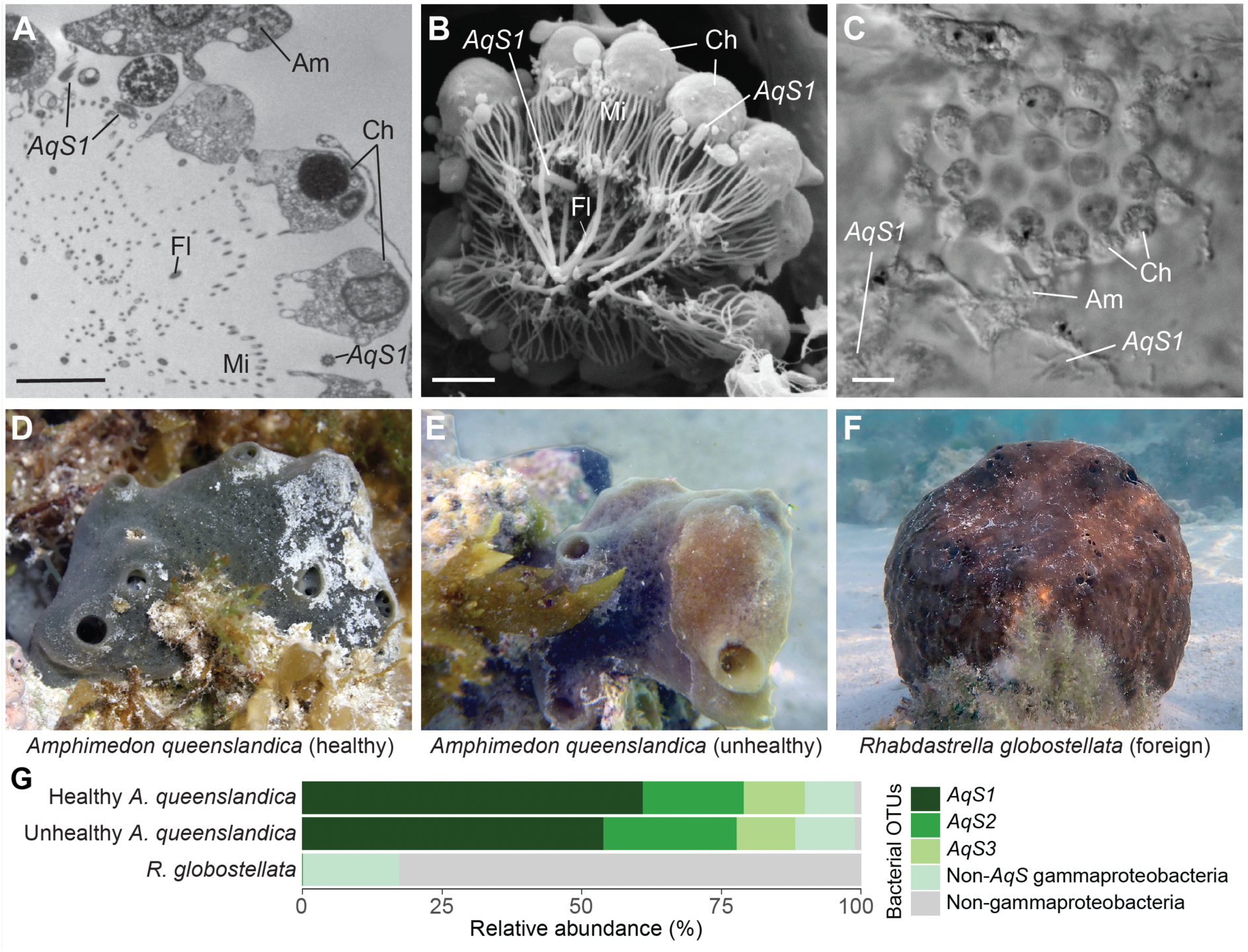
Native symbionts of *Amphimedon queenslandica* and contrasting bacterial communities of a co-occurring sponge *Rhabdastrella globostellata*. (A-C) Localisation of the dominant gammaproteobacterial symbiont *AqS1* near choanocyte chambers and within the mesohyl of juvenile and adult *A. queenslandica*: (A) Transmission electron micrograph showing choanocytes (Ch) with their microvillar collars (Mi) and flagella (Fl), a proximal amoebocyte (Am), and *AqS1* in the vicinity of the chamber. (B) Freeze–fracture scanning electron micrograph of choanocyte chamber with associated *AqS1*. (C) Differential interference contrast microscopy image showing motile amoebocytes and *AqS1* clusters adjacent to a juvenile choanocyte chamber. Images acquired using methods as in [28]. Scale bars: (A-C) 5 µm. (D–F) In situ morphology of sponges used for bacterial enrichments: (D) Healthy *A. queenslandica,* (E) unhealthy *A. queenslandica* showing reduced cellular density and pale colouration, and (F) *R. globostellata.* (G) Relative abundance of native symbionts *AqS1*, *AqS2* and *AqS3* and other bacterial OTUs in healthy and unhealthy *A. queenslandica* and in *R. globostellata* based on 16S rRNA gene profiling (see Table S1 for full OTU lists). *AqS1–3* dominate healthy *A. queenslandica* but are reduced in unhealthy individuals and absent from *R. globostellata*. These anatomical and microbiome differences provide the framework for comparing early immune regulatory responses to native versus foreign bacteria.

In the marine sponge *Amphimedon queenslandica*, bacterial symbionts are vertically transmitted during embryogenesis via maternal provisioning; however, immune discrimination is not confined to early development. We previously showed that larval settlement and metamorphosis disrupt the vertically inherited microbiome and coincide with the entry of environmental microbes into the developing juvenile [28]. This transition represents a critical life-history stage at which the host must actively discriminate resident symbionts from newly encountered bacteria to re-establish a stable microbiome. Thus, while vertical transmission establishes initial associations with high fidelity, post-metamorphic immune regulation likely contributes to symbiont maintenance and community resilience throughout juvenile and adult life.

Here, we use *A. queenslandica* to identify the earliest detectable cellular and transcriptional responses associated with discrimination between symbiotic and non-symbiotic bacteria during this post-metamorphic phase. We exposed actively feeding juvenile sponges, which already harbour vertically-inherited symbionts near choanocyte chambers and within the mesohyl (Fig. 1A-C), to native symbiotic bacteria and foreign bacterial communities via the aquiferous system. This feeding-based approach provides a tractable and ecologically relevant means to compare early host responses under controlled conditions, while enrichment of native symbionts amplifies early regulatory signals that might otherwise be difficult to resolve. Focusing on TF activation and nuclear translocation as a mechanistically informative readout of immune activity, we show that symbiotic bacteria are processed more rapidly and trigger early, transient activation and nuclear translocation of conserved innate immune TFs, whereas foreign bacteria are processed more slowly and elicit a distinct xenobiotic transcriptional response.

## MATERIALS AND METHODS

### Enrichment and characterisation of sponge bacterial communities

*Amphimedon queenslandica* and *Rhabdastrella globostellata* were collected from Heron Reef, Great Barrier Reef, Australia (23◦27ʹS, 151◦55ʹE) (Fig. 1D, E, F), and bacterial communities were enriched from individual sponges as previously described [30, 31]. The health of *A. queenslandica* individuals was visually assessed in the field prior to collection. Healthy individuals were dark blue-grey with high cellular mass, whereas unhealthy individuals had pale brown to olive regions with reduced cell mass (Fig. 1D, E). Bacterial enrichments from *A. queenslandica* and *R. globostellata* were diluted in 0.2 µm filtered sea water (FSW) to ∼ 10^7^ cells/ml. DNA was extracted from ∼ 10^6^ cells per enrichment for 16S rRNA gene amplicon sequencing to identify prokaryotic operational taxonomic units (OTUs) as previously described [28]. 16S profiling confirmed that *AqS1–3* dominated enrichments from *A. queenslandica* and were absent from *R. globostellata* enrichments, validating their use as native and foreign bacterial treatments, respectively (Table S1).

All field collections were conducted under Great Barrier Reef Marine Park Authority Permit G16/38120.1 issued to BM Degnan and SM Degnan.

### Rearing *A. queenslandica* larvae and juveniles

Naturally released *A. queenslandica* larvae were collected in larval traps from 25 individuals, pooled, and maintained in ambient seawater for 6-8 h before being induced to settle on the coralline alga *Amphiroa fragilissima* as previously described [32, 33] Within 6 h of settlement, postlarvae were detached from the algae and resettled individually onto glass coverslips [34] . Coverslips were placed in sterile 24-well plates containing 2 ml FSW and postlarvae were reared for 96 h at 25°C until reaching the juvenile stage with a functional aquiferous system, determined by the presence of a visible osculum. Juveniles at this stage actively filter-feed and represent the post-metamorphic phase in which environmental microbes first enter the sponge body via feeding [35].

### Labelling and tracing bacteria in juvenile sponges

Bacterial enrichments were centrifuged at 5000 x g for 10 min and resuspended in 1 ml of 5 µM carboxyfluorescein diacetate succinimidyl ester (CFDA-SE; Vybrant, Thermofisher Scientific) in FSW. After 15 min at room temperature, suspensions were centrifuged at 5000 x g for 10 min; pellets were washed twice with FSW, resuspended in 1 ml FSW, and maintained at 4°C in the dark. Juveniles were washed in FSW and the volume of FSW in each well was adjusted to 1.5 ml. CFDA-SE labelled bacterial enrichments from healthy *A. queenslandica*, unhealthy *A. queenslandica,* or *R. globostellata* (40 µl per well) were added to achieve similar final bacterial concentrations across treatments (2.7–3.6 × 10^4^ cells/ml), minimizing dose-dependent effects. Control juveniles received 40 µl FSW only, with no bacteria. Juveniles were incubated with labelled bacteria in the dark, and fixed at 0.5, 1, 2 and 8 h post-exposure (hpe) as previously described [36]. For 8 hpe incubations, wells were washed once with fresh FSW after 2 h and juveniles were reared in FSW only for the last 6 h. Fixed juveniles were stained with 4ʹ,6-diamidino-2-phenylindole (DAPI; 1:1000, Molecular Probes), mounted in ProLong Gold anti-fade (Molecular Probes) and imaged on a Zeiss LSM 510 META confocal microscope [37]. Labeled bacteria were readily internalized by sponge cells across all treatments.

For quantification, numbers of choanocyte chambers, archaeocytes and bacteria-containing archaeocytes were counted in four randomly-chosen confocal sections from the inner cortex of five juveniles per treatment per timepoint (n = 20 sections per juvenile). Archaeocytes were identified based on large size, amoeboid morphology, mesohyl localisation, phagocytic inclusions, and a conspicuous nucleolus [26, 34]. Counts from multiple confocal sections were averaged per individual juvenile, and juveniles were treated as independent biological replicates. Differences among treatments were tested using one-way ANOVA followed by Tukey’s HSD.

### CEL-seq2 analysis of juvenile transcriptomes

Individual juveniles were exposed to FSW (no bacteria controls), or bacterial enrichments from healthy *A. queenslandica* (healthy native), unhealthy *A. queenslandica* (unhealthy native), or *R. globostellata* (foreign) for 0, 2 or 8 h as described above. Each treatment and time point comprised four or five independent biological replicates, each representing a single individual juvenile (Table S2). At 0, 2 and 8 hpe, juveniles were transferred individually to 300 µl of RNALater (Sigma), stored overnight at 4°C, then at -20°C. RNA was extracted using the EZ Spin Column Total RNA Isolation Kit (BioBasic Inc., Toronto, Canada) according to the manufacturer’s instructions, and amplified and sequenced using CEL-seq2 [38]. Samples were multiplexed into two libraries and sequenced across two Illumina HiSeq2500 runs, with treatments and time points balanced across runs (Table S2). Reads were processed using the CEL-seq2 pipeline (available at https://github.com/yanailab/CEL-Seq-pipeline), then demultiplexed raw reads were mapped to the *A. queenslandica* genome and gene-level counts generated using Aqu 2.1 gene models [39] (Table S2). Data are available at NCBI Bioproject PRJNA1121035.

### Analysis of gene expression

Differential expression analysis used DESeq2 [40]. Prior to analysis, genes with median read counts < 10 were removed and filtered counts were normalised using variance stabilizing transformation (VST) [40]. An unsupervised principal component analysis (PCA) was visualised using ggplot2 [41]. Differentially expressed genes (adjusted *p* < 0.05) were identified by pairwise comparisons of each treatment against its matched FSW control. An UpSet plot (ComplexUpset and ggplot2) was used to visualise intersections [41, 42], and heatmaps (ComplexHeatmap) using normalised VST counts were used to visualise relative gene expression changes [43]. Genes were annotated as transcription factors (TFs), pattern recognition receptors (PRRs), or pathway components using curated annotations and eggNOG-mapper functional assignments.

Sparse partial least squares discriminant analysis (sPLS-DA; mixOmics) [44] was used to identify feature genes. A predictive model was trained on 2 hpe datasets using repeating 3-fold cross-validation (50 repeats), then used to predict bacterial type for 8 hpe bacterial datasets; only correctly predicted datasets were retained. sPLS-DA was then performed on the combined datasets (retained 8 hpe plus all 2 hpe) to identify feature genes by principal component. Samples were visualised by scatter plot with 95% confidence ellipses (ggplot2), and heatmaps were generated from VST counts (ComplexHeatmap).

A weighted co-expression analysis (WGCNA) [45] was performed on VST counts filtered by expression and variance (8457 genes with median or median absolute deviation in top 5000 were retained), using a signed Nowick-type topological overlap metric with soft threshold of β = 3. Co-expression modules were defined using dynamic tree cut (height 0.15; minimum module size 100). Module eigengenes were correlated with treatments and time points; modules with *p* < 0.05 were considered significantly associated with groups. Visualisation used ggplot2 and ComplexHeatmap [41, 43].

### GO and KEGG Pathway enrichment analyses

Gene ontology (GO) and Kyoto Encyclopedia of Genes and Genomes (KEGG) terms were assigned to the predicted proteome of *A*. *queenslandica* using eggNOG-mapper v2 [46]. KEGG enrichment for each module used one-sided Fisher’s exact tests in clusterProfiler [47] with FDR correction (*q* < 0.05). Distributions of significantly enriched terms across modules were visualised using ggplot2 heatmaps [41].

### Immunofluorescence

Juveniles were fixed at 1 and 2 hpe, and immunolabelled using rabbit polyclonal antisera raised against *A. queenslandica* IRF (TRSGSSADEQEPERPER), STAT (QRLFNHEDDPNRNE) and NF-κB (NGIDPSSLPEALIR) peptide epitopes (GenScript). Antisera were diluted 1:500 in blocking buffer and incubated with fixed juveniles overnight at 4°C as previously described [48]. Specificity controls included pre-immune sera from the same rabbits tested under identical conditions, which produced no detectable signal (Fig. S1). The secondary antibody (Alexa Fluor 647 goat anti-rabbit IgG) does not produce non-specific immunofluorescence in *A*. *queenslandica* [37, 49]. Nuclear localisation of TFs was assessed relative to Hoechst 33342 staining, and imaging settings were held constant across treatments.

To test whether heat-inactivation of native bacteria affected TF localisation, a healthy native bacterial enrichment (see above) was diluted in FSW to ∼ 10^7^ cells/ml, divided in half, and either maintained in the dark at 4°C or heated to 85°C for 15 min. Juveniles were exposed to 40 µl of either unheated or heat-inactivated native bacterial preparation as described above, fixed at 2 hpe, and immunolabelled with anti-IRF, anti-NF-κB or anti-STAT antisera as described above.

## RESULTS

To identify the earliest detectable immune discrimination between symbiotic and non-symbiotic bacteria during post-metamorphic life, we examined cellular processing and TF activation following bacterial uptake in juvenile *Amphimedon queenslandica*. Because TF activation and nuclear translocation initiate immune effector gene expression, we used these events as a readout of early immune regulatory commitment downstream of bacterial capture. Here, “early” refers to the first detectable regulatory divergence after bacterial uptake, rather than the moment of physical capture. We compared responses to native symbiotic and foreign bacterial communities introduced via filter-feeding against FSW controls. All analyses were performed on individual juveniles (n = 4 or 5 per treatment per time point; Table S2); replicates were derived from independent juveniles originating from pooled larval cohorts.

### *Amphimedon queenslandica* hosts a simple symbiosis that differs from a co-occurring sponge

Marine sponges typically host species-specific bacterial communities that differ from surrounding seawater and range from high to low complexity and abundance [10, 50–52]. *A. queenslandica* (Fig. 1D, E) has a low-complexity microbiome dominated throughout its life cycle by three vertically transmitted gammaproteobacterial symbionts (hereafter *AqS1*, *AqS2*, and *AqS3*) that together can comprise up to 90% of the bacterial community (Fig. 1G) [28, 30]. Unhealthy individuals, identified by reduced cellular density and paler colouration (Fig. 1E), showed reduced relative abundance of these core symbionts compared with healthy individuals, consistent with early dysbiosis (Fig. 1F and Table S1). In contrast, the co-occurring sponge *Rhabdastrella globostellata* hosts a distinct and more complex bacterial community that does not include *AqS1–3* (Fig. 1F, G and Table S1). These contrasting microbiomes provided a framework for comparing host responses to symbiotic (native) versus foreign bacterial communities.

### Native symbionts are rapidly transported from choanocytes to internal phagocytic cells

Sponges possess an internal epithelial boundary that includes choanocyte chambers and an internal collagenous mesohyl populated by phagocytic, secretory, biomineralizing, and stem cell types that interact with both food bacteria and symbionts [24–27]. To examine early cellular handling, we introduced CFDA-SE–labelled bacterial communities enriched from healthy *A. queenslandica* (healthy native), unhealthy *A. queenslandica* (unhealthy native) or *R. globostellata* (foreign) into filtered seawater surrounding juvenile *A. queenslandica* (Fig. 2A).

**Fig. 2.**
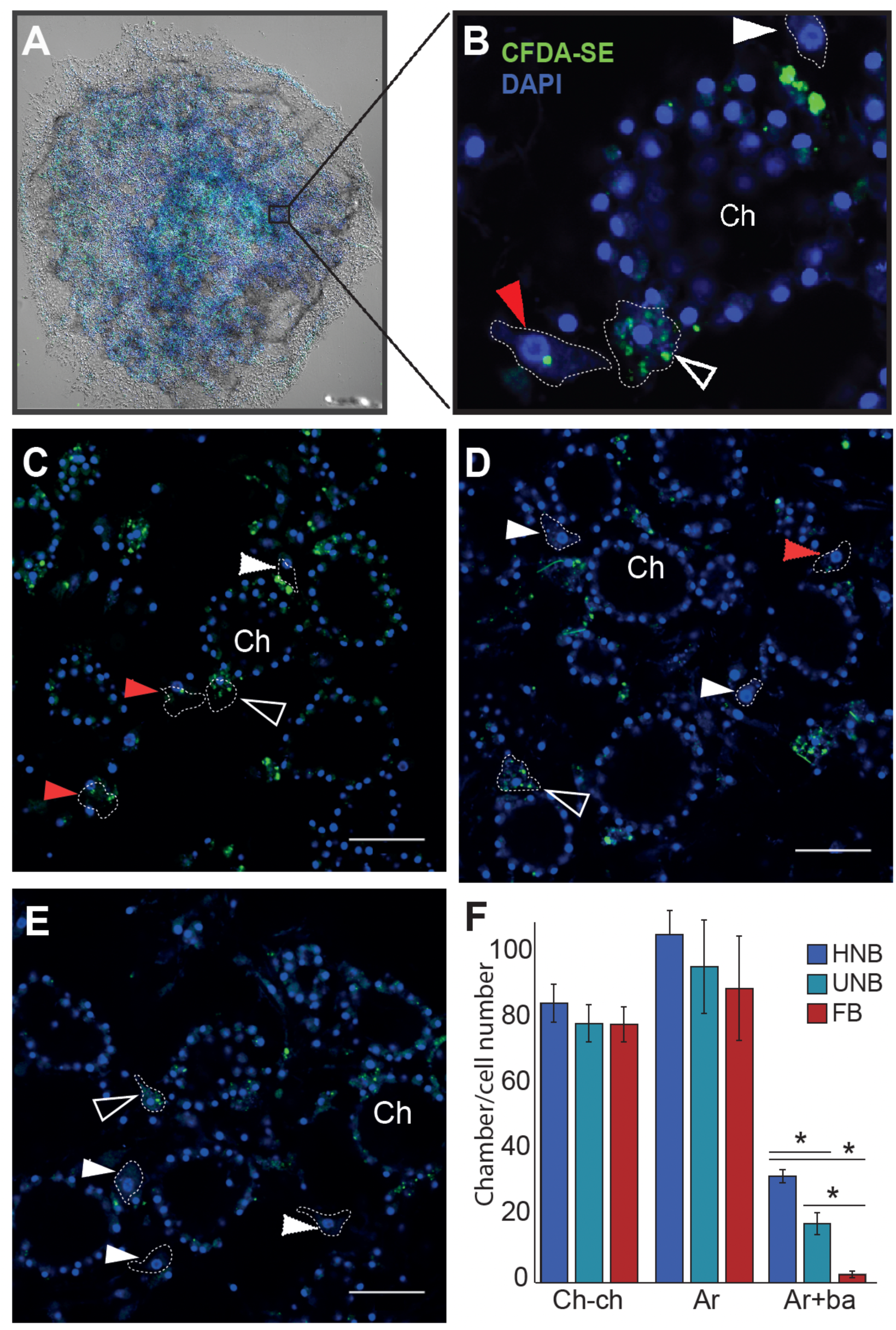
Differential cellular processing of native and foreign bacteria in juvenile *Amphimedon. queenslandica*. (A) DIC and fluorescence image of a DAPI-stained (blue) juvenile exposed to CFDA-SE-labelled (green) native bacteria at 8 h post-exposure. (B) Confocal micrograph of a choanocyte chamber (Ch) showing labelled microbes within choanocytes and adjacent amoeboid cells; a bacteria-containing amoebocyte (open arrowhead), an bacteria-containing archaeocyte with prominent nucleolus (red arrowhead), and an archaeocyte lacking bacteria (white arrowhead) are outlined. (C-E) Fluorescence micrographs of juveniles at 2 hpe following exposure to bacteria from healthy native (HNB), unhealthy native (UNB), or foreign (FB; *Rhabdastrella globostellata*) sources, respectively. Microbes are detected in choanocyte chambers and proximal amoebocytes in all three treatments. Scale bars, 25 µm. (F) Quantification of choanocyte chambers (Ch-ch), archaeocytes (Ar) and bacteria-containing archaeocytes (Ar+ba) at 2 hpe (mean ± s.e.m.). Native bacteria are transferred more frequently to archaeocytes than foreign bacteria (\**p* < 0.01; Tukey’s HSD).

Within 30 min, both native and foreign bacteria were detected within choanocytes and in adjacent amoebocytes contacting the basal surface of choanocytes (Fig. 2B and Fig. S2), indicating broadly similar early capture and initial association across treatments. By 2 hours post exposure (2 hpe), however, downstream cellular processing diverged. Native bacteria (healthy and unhealthy) were efficiently transferred to archaeocytes, which are large phagocytic mesohyl cells with amoeboid morphology, whereas foreign bacteria were rarely detected in archaeocytes (p < 0.01; Fig. 2C-F and Fig. S3). Moreover, archaeocytes in healthy native treatments contained significantly more bacteria that in unhealthy native treatments (Fig. 2F), indicating that downstream handling differs not only between native and foreign bacteria but also between native communities differing in state.

### Native symbionts trigger a rapid, transient transcriptional response distinct from the foreign bacteria response

To determine when discrimination becomes detectable at the molecular level, we compared transcriptomes of juveniles exposed to healthy native, unhealthy native, or foreign bacterial communities for 2 and 8 hours with matched filtered sea water (FSW) controls (Table S2). Principal component analysis revealed distinct trajectories, with native and foreign treatments separating strongly along PC1 (Fig. 3A).

**Fig. 3.**
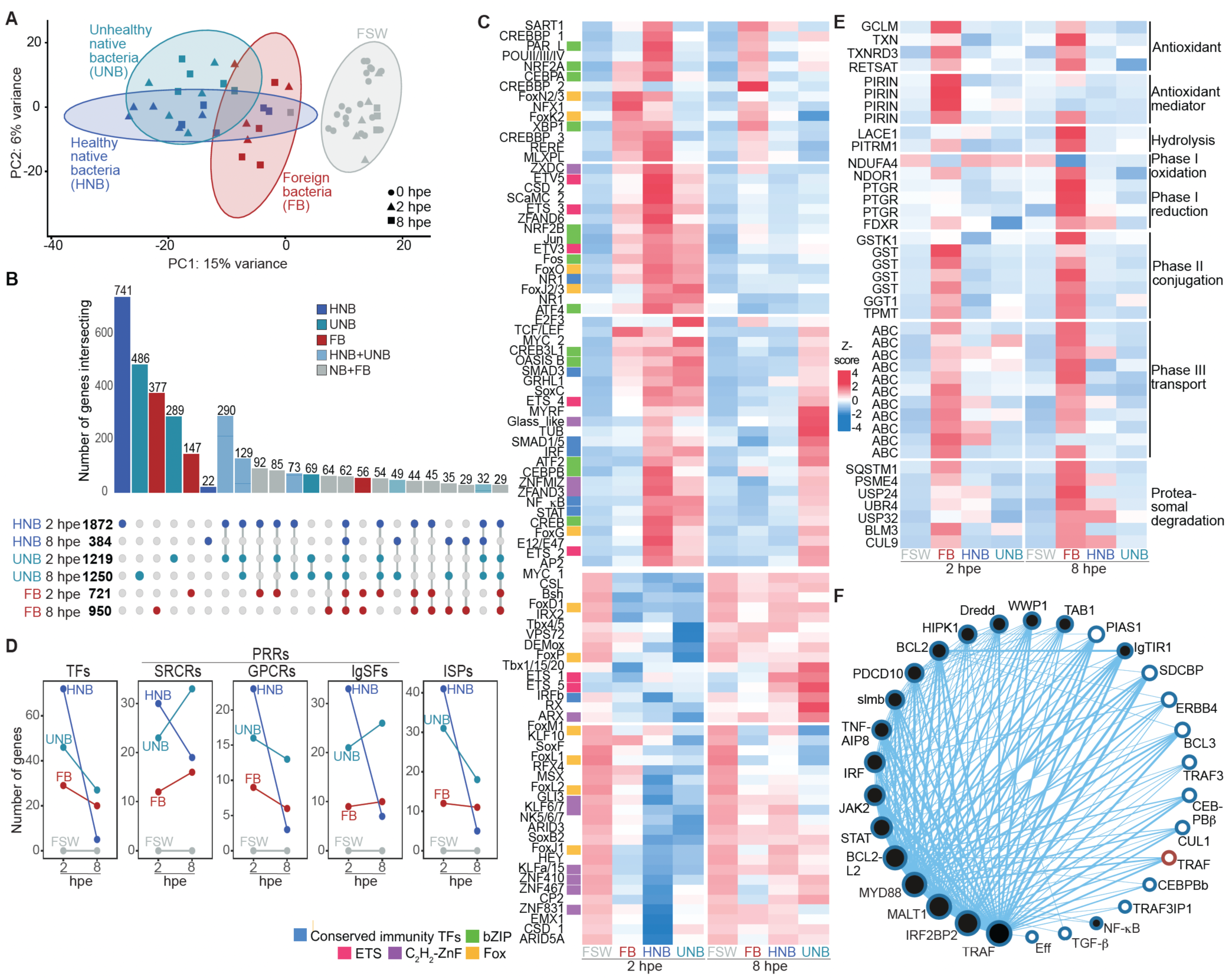
Native symbionts elicit a rapid, transient immune transcriptional program, whereas foreign bacteria induce xenobiotic pathways symbionts. (A) Principal component analysis (PCA) of juvenile *A. queenslandica* transcriptomes following exposure to filtered seawater (FSW, grey) or bacterial enrichments from *R. globostellata* (foreign bacteria; FB, red); or from healthy (HNB, blue) and unhealthy (UNB, cyan) *A. queenslandica.* Symbols denote timepoints (0 hpe circles; 2 hpe triangles; 8 hpe squares); ellipses show 95% confidence intervals. (B) UpSet plot summarising differentially expressed genes (DESeq2; *p*-adj < 0.05) in each bacterial treatment at 2 and 8 hpe relative to time-matched FSW controls. The number of differentially expressed genes are shown at left in bold; intersections between treatments are shown above bars. The colour coding follows (A) except for intersections between healthy and unhealthy native treatment (teal), and between native and foreign treatments (grey). (C) Hierarchically clustered heatmap of differentially expressed transcription factor (TF) genes across treatments and timepoints (z-scored expression shown). Order of samples is shown below heatmap, and TF names are shown at left. Conserved immunity (blue), bZIP (green), ETS (red), C2H2 zinc-finger (purple) and Fox (yellow) TFs are marked by an adjacent box. (D) Slope charts of numbers of differentially expressed TFs, pattern recognition receptors (PRRs; SRCRs, GPCRs, IgSFs) and immune signalling pathway components (ISPs) at 2 and 8 hpe for each bacterial treatment relative to FSW controls. See Fig. S4 for heatmap of differentially expressed PRRs and ISPs). (E) Heatmap pf xenobiotic metabolism-associated genes activated by foreign bacteria, annotated by functional phase (I-III) and related processes (z-scored expression shown; sample order as in C). (F) Co-expression network showing the top 30 most connected genes in the healthy native bacteria-induced response. Node size reflects number of connections. Black-filled nodes indicate genes also identified as discriminating features by sPLS-DA (Fig. S7; Table S5). All genes are in WGCNA module 1 (blue), except for TRAF in module 2 (brown) (Fig. S8; Tables S6, S7).

Healthy native bacteria elicited a rapid and transient transcriptional response relative to FSW controls. Approximately twice as many genes were differentially expressed at 2 hpe compared with juveniles exposed to foreign bacteria (DESeq2 adjusted p < 0.05; Fig. 3B–E, S3 Fig. S4; Table S3). By 8 hpe, expression in native-exposed juveniles largely returned to FSW baseline, indicating tight temporal regulation. Unhealthy native bacteria induced qualitatively similar but attenuated responses, with delayed kinetics that paralleled the slower transfer to archaeocytes (Fig. 2F).

In contrast, foreign bacteria induced a weaker early response but a more prolonged trajectory, involving a distinct gene set that remained divergent at 8 hpe (Fig. 3A–E). Genes uniquely enriched in the foreign-bacteria response included those involved in xenobiotic metabolism and detoxification-associated pathways (Fig. 3E, Fig. S4 and Table S3), indicating engagement of a downstream program distinct from the transient immune activation induced by native symbionts. Together, these graded responses suggest that transcriptional regulation mirrors early cellular handling of bacteria.

### Early discrimination is marked by coordinated, transient activation of immune transcription factors

Among genes induced by native symbionts, transcription factors were prominently enriched. At 2 hpe, more than 80 TFs were significantly upregulated (p-adj < 0.05), with most returning to FSW baseline by 8 hpe (Fig. 3C). Transiently induced TFs included NF-κB, two IRFs, STAT, two SMADs, AP-1, three Fox genes, all ETS and 13 of 17 bZIP family members, and a nuclear receptor—many with established roles in bilaterian innate immunity [15, 21, 23, 53–58]. Adult cell-type expression profiles [34, 59] indicates that these TFs are largely constitutively expressed in epithelial choanocytes and pinacocytes but also present in mesohyl cell types (Fig. S5 and Table S4), consistent with immune readiness at the epithelial-mesohyl interface.

Responses to unhealthy native bacteria were delayed and weaker, with TF expression at 8 hpe resembling the 2 hpe response to healthy native bacteria (Fig. 3C). In contrast, foreign bacteria induced markedly fewer changes in immune TF expression at 2 hpe and lacked the coordinated, transient TF activation characteristic of native symbionts (Fig. 3C), consistent with absence of coordinated TF nuclear translocation in amoebocytes exposed to foreign bacteria (see below).

### Native bacteria induce transient immune receptor and pathway activation, whereas foreign bacteria elicit xenobiotic-associated programs

Expression of pattern recognition receptors (PRRs) broadly paralleled TF dynamics in response to native bacteria. Although expression changes do not imply direct receptor-level discrimination, differences between native and foreign bacterial treatments were observed. Healthy native bacteria induced transient differential expression of 56 scavenger receptor cysteine-rich (SRCR) genes, 32 G-protein coupled receptor (GPCR) genes, and 50 immunoglobulin superfamily genes relative to FSW controls (Fig. 3D, Fig. S4 and Table S3). Two Toll-like receptors (AmqIgTIR1 and AmqIgTIR2), comprising extracellular IL1R-like immunoglobulins and an intracellular TIR domain[16], were uniquely regulated by native bacteria. Several sponge-specific GPCRs with affinity to parahoxozoan γ-aminobutyric acid (GABA) receptors were also induced; GABA can be synthesized and degraded by *AqS1* and mediates host–microbe interactions in other holobionts [60–62].

Foreign bacteria induced a more limited PRR signature, including selective upregulation of multiple CL163-L1 SRCR genes (Fig. 3D, Fig. S4 and Table S3), a receptor class associated with inflammatory activation states in mammalian macrophages [63]. This distinct PRR signature occurred alongside weak activation of the native-associated immune TF program, consistent with foreign bacteria eliciting a qualitatively different downstream response rather than a scaled-down version of the native response.

Native symbionts also induced transient activation of multiple conserved innate immune pathways at 2 hpe, including eight TRAFs in the pro-inflammatory cytokine TNF pathway, and multiple components of Toll, Imd, apoptosis, TGF-β and JAK–STAT pathways (Fig. 3D, Fig. S4 and Table S3). Most of these pathway components are constitutively most highly expressed in adult epithelial pinacocytes and choanocytes (Figs S5, S6 and Table S4), consistent with an epithelium poised for rapid response. By contrast, foreign bacteria induced fewer innate immune pathway genes, generally at lower levels and with delayed kinetics, and instead elicited a transcriptional response dominated by xenobiotic metabolism pathways (Fig. 3E, Fig. S4 and Table S3). Xenobiotic-associated genes are chiefly constitutively most highly expressed in internal archaeocytes in adults Fig. S6 and Table S4), consistent with foreign bacteria engaging internal processing programs distinct from those induced by native symbionts.

### IRF, NF-κB, and STAT comprise a central regulatory module associated with early discrimination

To identify key regulators, we integrated DESeq2 results with sparse partial least squares discriminant analysis (sPLS-DA) and weighted gene co-expression network analysis (WGCNA). Two WGCNA modules were induced at 2 hpe by native bacteria; module 1 encompassed most immune genes identified by both DESeq2 and sPLS-DA (Fig. 3F, Figs S7, S8 and Tables S5, S6). The most highly connected genes within this module included orthologues of bilaterian innate immunity genes, with IRF, NF-κB, and STAT among the most connected TFs, together with genes involved in endocytosis and vesicle trafficking (Fig. 3F). Because TFs represent integrative nodes where multiple upstream signals converge, we next examined whether these TFs show differential nuclear translocation during early bacterial handling.

### TF nuclear translocation marks the earliest detectable regulatory divergence

Using custom antibodies, we tracked subcellular _ocalization of IRF, NF-κB, and STAT in juvenile *A. queenslandica* during the first two hours of exposure to FSW (control), healthy native bacteria or foreign bacteria (see Methods; Fig. 4 and Figs S1, S9, S10). In FSW controls, all three TFs were predominantly cytoplasmic in choanocytes and proximal amoebocytes, while NF-κB was also present in nuclei of a subset of archaeocytes near choanocyte chambers (Fig. 4A–C and Figs S9, S10).

**Fig. 4.**
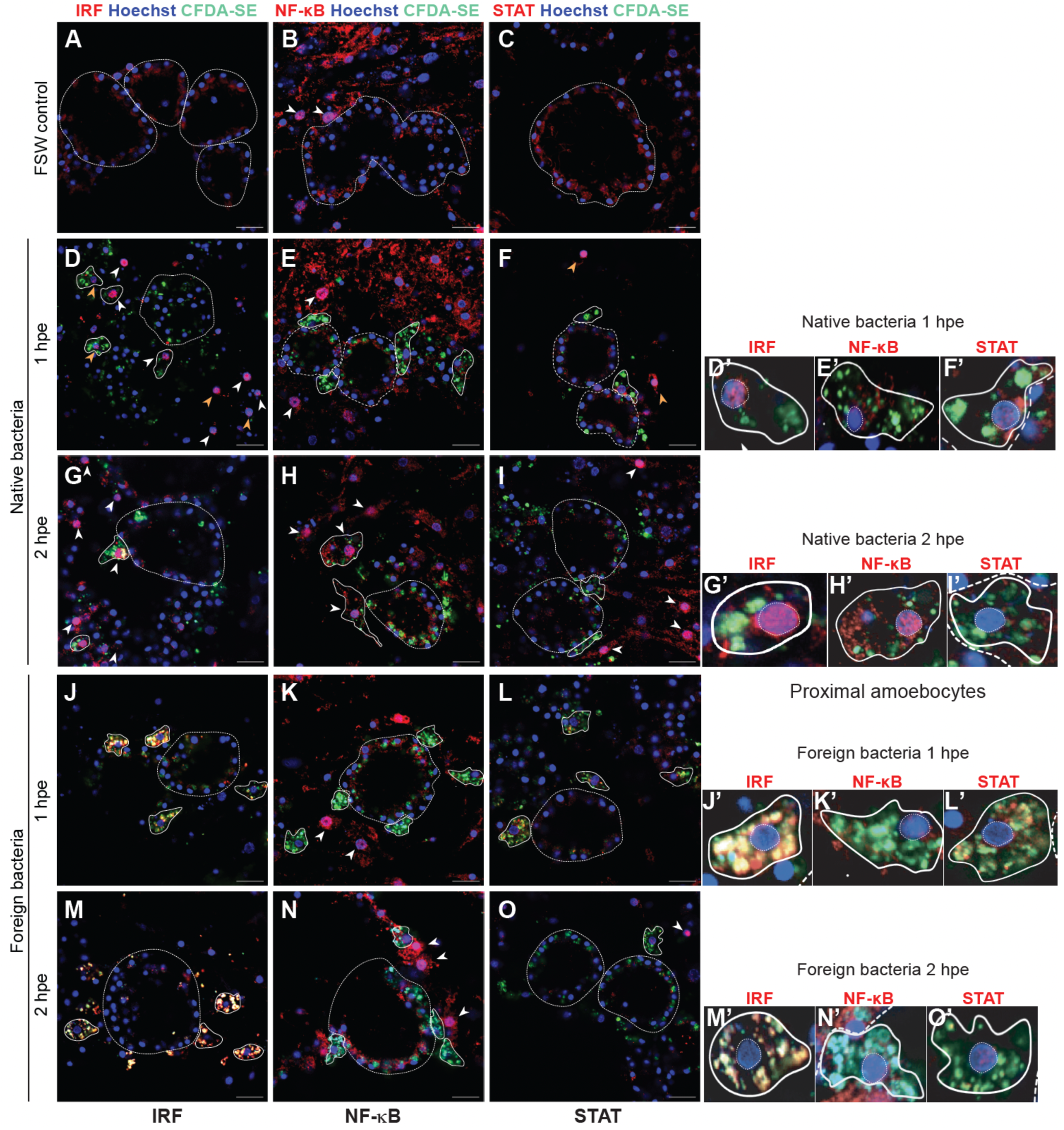
Subcellular localisation of IRF, NF-κB, and STAT during early responses to native and foreign bacteria. Immunofluorescence imaging of juvenile *Amphimedon queenslandica* exposed to filtered seawater (FSW control), native symbiotic bacteria, or foreign bacteria at 1 and 2 h post-exposure (hpe). Transcription factors are shown in red (IRF, NF-κB, or STAT), bacteria in green (CFDA-SE), and nuclei in blue (Hoechst). See Figs S9 and S10 for separated fluorescence channels. (A-C) FSW controls showing predominantly cytoplasmic localisation of all three TFs. (D-I) Healthy native bacteria induce rapid IRF nuclear translocation in proximal amoebocytes by 1 hpe (D), followed by NF-κB nuclear localisation at 2 hpe (H). STAT nuclear localisation occurs in a distinct subset of amoebocytes not directly associated with bacterial uptake (F, I). (J-O) Foreign bacteria do not induce coordinated nuclear localisation of IRF, NF-κB, or STAT at either time point; IRF signal remains cytoplasmic and frequently co-localises with internalised bacteria (J, M). Choanocyte chambers and proximal amoebocytes containing labelled bacteria are outlined with dashed and solid lines, respectively. White arrowheads indicate TF-positive amoebocyte nuclei; yellow arrowheads indicate TF-positive nuclei in other amoebocytes. Scale bars, 10 µm. Panels with primes (ʹ) show magnified views of representative proximal amoebocytes with nuclei outlined. Nuclear localisation was scored based on overlap with Hoechst signal.

Within one hour of exposure to native bacteria, IRF translocated to nuclei of proximal amoebocytes that had engulfed these bacteria (Fig. 4D). In contrast, in foreign-exposed juveniles, IRF remained cytoplasmic in proximal amoebocytes and co-localised with engulfed bacteria (Fig. 4J). At this time, NF-κB and STAT did not localise to nuclei of amoebocytes directly engaged in bacterial uptake in either treatment (Fig. 4E, F, K, L). Native bacteria did, however, induce nuclear localisation of STAT in a distinct subset of amoebocytes not directly associated with bacterial uptake and more distant from choanocyte chambers (Fig. 4F), consistent with native bacteria indirectly inducing STAT nuclear localisation. By 2 hpe, NF-κB co-localised with IRF in nuclei of amoebocytes that had engulfed native bacteria but were no longer in direct contact with choanocyte chambers (Fig. 4G, H) consistent with sequential activation. STAT nuclear localisation remained restricted to native bacteria-exposed juveniles in amoebocytes not directly associated with bacterial uptake (Fig. 4I).

Together, these data indicate that differential IRF behaviour in proximal amoebocytes within the first hour is the earliest detectable regulatory divergence between symbiotic and non-symbiotic bacteria in *A. queenslandica*. The nuclear localisation of STAT at the same time in distant amoebocytes appeared to be induced by an intercellular signal rather than by direct interaction with native bacteria. Both responses required a signal from living native bacteria, as exposure to heat-killed native bacteria did not induce nuclear translocation of either IRF or STAT (Fig. S11).

## Discussion

Juvenile *Amphimedon queenslandica* respond to native (symbiotic) and foreign bacteria with markedly different kinetics and regulatory programs, demonstrating that innate immune regulation is central to sponge–microbe interactions. Native bacteria rapidly engage a broad, transient transcriptional response enriched for innate immune pathways and TF families that are shared across animals, including IRF, STAT, NF-κB, AP-1 and other bZIPs, ETS family members, SMADs and nuclear receptors [15, 21, 23, 53–58]. Although effector functions cannot yet be directly assayed in *A. queenslandica,* TF activation and nuclear translocation represent a well-established commitment point in innate immunity, providing a mechanistically informative readout because these events initiate immune effector gene expression and shape response outcomes.

Importantly, our data identify the earliest detectable regulatory divergence following bacterial uptake, rather than the mechanisms of initial capture or cell-surface sensing. Choanocytes are the first cells to capture environmental bacteria, and multiple immune-related genes (including TFs and PRRs) are constitutively expressed in sponge epithelia. However, we do not determine whether discriminatory recognition occurs at the choanocyte surface, at other epithelial surfaces (e.g. endopinacoderm) or through specific receptors at initial capture. Instead, our findings show that discrimination becomes evident downstream of capture, where TF nuclear translocation occurs primarily in amoebocytes during early bacterial processing.

Responses to unhealthy native bacteria were delayed and attenuated, mirroring slower cellular transfer and supporting the idea that sponge innate immunity can tune regulatory responses o changes in microbial community state. This graded response suggests that TF-mediated regulation enables modulation of host programs in response to subtle shifts consistent with the onset of dysbiosis. While we cannot yet resolve whether these responses promote symbiont retention, clearance, or rebalancing, their timing and magnitude indicate that transcriptional control is a key component of early sponge–microbe interactions.

In contrast to native bacteria, foreign bacteria elicited a comparatively weak activation of innate immune programs and instead induced a transcriptional program dominated by xenobiotic metabolic processes. This indicates that foreign bacteria are processed through a distinct downstream program, consistent with foreign microbes presenting primarily a chemical or metabolic challenge and triggering detoxification or degradation responses rather than strong engagement of canonical immune TF cascades [64, 65]. Consistent with this interpretation, IRF and NF-κB did not translocate to nuclei in amoebocytes containing foreign bacteria at 2 hpe, indicating that foreign bacteria do not engage the coordinated TF-mediated immune module activated by native symbionts.

Sponges lack a gut, nervous system and muscle, but do have an epithelium and underlying phagocytic mesohyl cell types comparable – and in some cases arguably homologous – to those of other animals [15, 25, 27]. The constitutive expression of immune TFs, PRRs and signalling components in sponge epithelia, together with rapid activation and nuclear translocation of TFs in adjacent amoebocytes, supports a model in which immune readiness is embedded within the epithelial–mesohyl interface.

Given the close association between symbionts and epithelia across disparate animals [1, 3, 6, 66], epithelial boundaries likely represent conserved sites where early immune regulatory processes contribute to discrimination among beneficial, harmful and neutral microbes.

Although PRR expression patterns differed between native and foreign exposures, our data do not establish a causal role for specific receptors in mediating discrimination. These PRR expression changes more likely reflect downstream regulatory programs coordinated by TF activation than receptor-level specificity at initial contact.

IRF emerged as the earliest discriminating regulator in this system. IRF rapidly translocated to nuclei of amoebocytes that had engulfed native bacteria, but remained cytoplasmic and co-localised with engulfed foreign bacteria. This behaviour marks the first clear regulatory divergence observed in our study. IRF co-localisation with foreign bacteria is consistent with observations in mammals where IRF can participate in endosomal signalling through adaptor proteins such as MyD88 [55, 67–69], raising the possibility that IRF integrates bacterial context to determine whether transcriptional activation is initiated. NF-κB translocated to nuclei shortly after IRF in symbiont-containing amoebocytes, consistent with activation, whereas STAT nuclear localisation occurred in a distinct subset of amoebocytes not directly engaged in bacterial uptake, consistent with secondary intercellular signalling via pathways such as JAK–STAT, Toll, Imd, Wnt, FoxO, and TGF-β. Amoebocytes share functional similarities with macrophages in other animals and participate in sponge defense [27, 59]; however, we do not directly test their developmental or evolutionary homology here. Nonetheless, our data indicate that discrimination is coordinated across multiple cell types during early processing rather than restricted to the site of bacterial uptake.

By modelling the post-metamorphic phase, when environmental microbes begin entering the juvenile and the vertically inherited microbiome is restructured, our feeding-based experimental design targets immune mechanisms involved in symbiont maintenance and regulation through the sponge life cycle. Within this framework, we resolve the earliest detectable regulatory divergence that follows bacterial uptake: native symbionts rapidly engage a coordinated innate immune module centred on TF activation and nuclear translocation in amoebocytes, whereas foreign bacteria elicit a xenobiotic-dominated transcriptional program and do not engage the same TF module. This divergence becomes evident shortly after uptake, and downstream of initial capture by filter-feeding, identifying TF activation as an early regulatory checkpoint in sponge–microbe interactions. Together, these findings provide a mechanistic basis for differential processing of microbial partners in aquatic filter-feeding animals that face continual environmental exposure.

## Acknowledgments

The authors thank staff at the Heron Island Research Station for logistical assistance during field collections, and Nick Rhodes at QCIF for management of high-performance computing facilities.

## Author contributions

Conceptualization: SMD, BMD Methodology: SMD, BMD, BYang, BYuen, HY Investigation: BYang, BYuen, HY, SMD, BMD

Visualization: BYang, BYuen

Funding acquisition: SMD, BMD Project administration: SMD, BMD Supervision: SMD, BMD

Writing – original draft: SMD, BMD, BYang Writing – review & editing: SMD, BMD, BYang, HY

## Data and materials availability

Transcriptomes generated by CelSeq2 as part of this study are available at NCBI Bioproject PRJNA1121035. All other data are available in the main text or the Supporting Information.

