## Supplementary Information for "Early transcription factor activation distinguishes symbiotic from non-symbiotic bacteria during microbiome processing in a sponge"

**Supplementary Figure Legends**

**Fig. S1. Pre-immune sera do not cross react with juvenile *A. queenslandica*.**
Immunostaining performed with pre-immune sera pooled from three rabbits before they were individually injected with IRF, STAT or NF-κB peptides showed no cross reactivity to juvenile *A. queenslandica*. No cross reactivity was also observed when individual pre-immune sera were applied.
(A-C) Pre-immune sera plus goat anti-rabbit IgG (H+L) secondary antibody (Alexa Fluor 647; red) in juveniles exposed to (A) filtered seawater, (B) foreign bacteria or (C) native bacteria. Showing little to no detectable background. Previous analyses confirmed that the goat anti-rabbit IgG (H+L) secondary antibody (Alexa Fluor 647) also does not generate background fluorescence in juveniles [1, 2]. Choanocyte chambers outlined with dotted lines.
(A'-C') CFDA-SE-labelled bacterial cells (green). Mesohyl cells in the vicinity of choanocyte chambers with a distinct bacterial load are outlined by solid lines.
(A''-C'') Hoechst-stained nuclei (blue).
(A'''-C''') Merged images showing immunostaining background and introduced bacterial cells in juvenile cells.
Scale bars, 10 µm.

**Fig. S2. Uptake and fate of labelled native and foreign microbes in juvenile *A. queenslandica* at 0.5 and 1 hpe.**
(A-C) DAPI-stained juveniles (blue nuclei) 0.5 hpe to microbes labelled with CFDA-SE (green).
(D-F) DAPI-stained juveniles 1.0 hpe to microbes labelled with CFDA-SE. A single representative choanocyte chamber (Ch) is labelled in each micrograph.
(A, D) Juveniles exposed to microbes from healthy *A. queenslandica*.
(B, E) Juveniles exposed to microbes from unhealthy *A. queenslandica*.
(C, F) Juveniles exposed to microbes from *R. globostellata*.
Open arrowheads point to representative amoebocytes adjacent to a choanocyte chamber containing labelled microbes. White arrowheads point to representative archaeocytes without microbes.
Scale bars, 25 µm.

**Fig. S3. Uptake and fate of labelled native and foreign microbes in juvenile *A. queenslandica* at 8 hpe.**
DAPI-stained juveniles (blue nuclei) exposed to bacteria labelled with CFDA-SE (green).
(A, C, E) Confocal fluorescence micrographs.
(B, D, F) Differential interference contrast micrographs.
(A, B) Juveniles exposed to microbes from healthy *A. queenslandica*.
(C, D) Juveniles exposed to microbes from unhealthy *A. queenslandica*.
(E, F) Juveniles exposed to microbes from *R. globostellata*.

A single representative choanocyte chamber (Ch) is labelled in each micrograph. Red arrowheads point to representative archaeocytes with microbes.
Scale bars, 25 µm.

**Fig. S4. Immediate transcriptional response of innate immunity genes to native and foreign bacteria.**
Hierarchically clustered heatmaps of differentially expressed (DESeq2 p-adj < 0.05).
(A-C) Expression of PPRs in response to native and foreign bacteria at 2 and 8 hpe. Subsets of PRRs are separated into (A) scavenger receptor cysteine-rich domain-containing receptors (SRCRs), (B) G-protein coupled receptors (GPCRs) and (C) immunoglobulin superfamily members (IgSFs).
(D) Innate immunity signalling pathways in response to native and foreign bacteria at 2 and 8 hpe. Gene names or IDs are shown to the left and pathway memberships to the right in D (S3 Table). z-score for both heat maps in the middle.

**Fig. S5. Adult cell type expression of immunity genes that are up- or down-regulated in response to native symbionts.**
(A, B) Expression of *A. queenslandica* TFs in adult cell types that (A) were manually isolated and sequenced [3], or (B) were separated in an unsupervised manner and sequenced [4].
(C, D) Expression of PRRs in adult cell types that (C) were manually isolated and sequenced, or (D) were separated in an unsupervised manner and sequenced.
(E) PCA plot of CEL-Seq2 transcriptomes of manually isolated cells with 95% confidence level ellipse plots. Blue, choanocytes; red, archaeocytes; green, pinacocytes. Adapted from [3].
(F) 2D projection of MARS-Seq transcriptomes of adult metacells and single cells generated by physically dissociating an adult sponge in calcium/magnesium-free seawater and generating single-cell libraries from random cells. Cell clusters with known or hypothesised identity are annotated and highlighted in grey. Adapted from [4].

**Fig. S6. Adult cell type expression of immunity and xenobiotic factors that are up- or down-regulated in response to native symbionts and foreign bacteria.**
(A, B) Expression of *A. queenslandica* immunity factors induced by native symbionts in adult cell types that (A) were manually isolated and sequenced [3], or (B) were separated in an unsupervised manner and sequenced [4].
(C, D) Expression of xenobiotic factor induced by foreign bacteria in adult cell types that (C) were manually isolated and sequenced, or (D) were separated in an unsupervised manner and sequenced.
(E) PCA plot of CEL-Seq2 transcriptomes of manually isolated cells with 95% confidence level ellipse plots. Blue, choanocytes; red, archaeocytes; green, pinacocytes. Adapted from [3].
(F) 2D projection of MARS-Seq transcriptomes of adult metacells and single cells generated by physically dissociating an adult sponge in calcium/magnesium-free seawater and generating single-cell libraries from random cells. Cell clusters with known or hypothesised identity are annotated and highlighted in grey. Adapted from [4].

**Fig. S7. sPLS-DA of juvenile transcriptional responses to native microbes from healthy and unhealthy *A. queenslandica*, and foreign microbes from *R. globostellata*.**

(A) sPLS-DA of the full 2 and 8 hpe datasets that passed the model checking (see Methods). 95% confidence ellipses shown.

(B) sPLS-DA plot showing 2 hpe transcriptomes involved in the model construction. These are linked by dashed lines: FSW (n=5); foreign microbes (FB; n=4); healthy-native bacteria (HNB; n=5); and unhealthy-native bacteria (UNB; n=5). 8 hpe transcriptomes participate in the prediction but not the model construction.

(C) Prediction results of sPLS-DA for 8 hpe transcriptomes based on the modelling of 2 hpe transcriptome features. The number of squares represents the number of samples; the triangle position indicates the predicted principal component.

(D, E) Hierarchically clustered heatmaps of the expression of the 380 and 90 protein coding genes selected from PC1 (D) and PC2 (E), respectively.

(F) KEGG pathway analysis of the 470 feature genes identified by the sPLS-DA, with 380 genes being more responsive to 77% of these are upregulated in juveniles exposed to native and foreign bacteria, respectively.

**Fig. S8. Co-expressed coding sequences in juveniles exposed to native and foreign bacteria using WGCNA.**

(A) Scaled boxplot of the relative expression profile for genes in co-expression modules 1-6 (Table S6). The same gene is connected by a solid line among treatment groups. FSW, filtered sea water control; FB, foreign bacterial treatment; HNB, healthy-native bacteria treatment; UNB, unhealthy-native bacterial treatment.

(B) Percentage of significantly differentially expressed coding sequences identified using DESeq2 ($p\text{-adj} < 0.05$) in each WGCNA co-expression module. The genes identified by DESeq2 as differentially expressed for native (NB) and foreign (FB) bacteria are divided into two sets, dark and light blue or red, respectively, depending on the expression profile. NDE, not significantly differentially expressed using DESeq2 ($p\text{-adj} > 0.05$).

(C) Relative changes in the eigengene of each co-expression module across treatment groups. The eigengene of each module was obtained by co-expression analysis, and the relative expression change was calculated by specifying its expression at 0 hpe to be zero. The number of coding sequences in each module shown in the top left corner.

(D) Heatmap of the correlations between the modules identified by WGCNA and each treatment group ($p\text{-value} < 0.05$ is indicated by *, and $p\text{-value} < 0.01$ is indicated by **).

(E) Distribution of 53 differentially expressed conserved immune pathway genes in co-expression modules.

(F) Scatter plot of the intramodular contributions of all genes in the native and foreign bacteria-associated modules 1 and 3, respectively, and their correlations with native and foreign bacterial traits identified in WGCNA (Table S7). The top 50% of genes in both module membership and gene significance are indicated by the dotted lines. Core genes in response to foreign bacteria identified in sPLS-DA are labelled with black fill.

(G) Network diagram of the top connected genes in the foreign bacteria-associated module 3. The connections between nodes are indicated by the edges, and the degree of connectivity is indicated

by the thickness of the edges. The number of edges is represented by the size of the nodes. Core genes in response to symbionts identified in sPLS-DA are labelled with black fill.
(H) Dotplot of KEGG pathway enrichment analysis on differentially upregulated genes that are uniquely induced by native or foreign bacteria at 2 hpf. The p-value obtained from Fisher's exact test for the respective pathway is represented by the color-coding scheme. The enriched gene ratio within the respective pathway is indicated by the size of a dot.

**Fig. S9. Immunolocalization of IRF, STAT, and NF-κB in response to native and foreign bacteria at 2 hpe.**
(A, D, G) Separated fluorescence channels of panels shown in Fig. 4 of 2 hpe juveniles exposed to FSW.
(B, E, H) Separated fluorescence channels of panels shown in Fig. 4 of 2 hpe juveniles exposed to foreign bacteria from *R. globostellata*.
(C, F, I) Separated fluorescence channels of panels shown in Fig. 4 of 2 hpe juveniles exposed to native bacteria from healthy *A. queenslandica*.
First column (letter, no prime), immunostained juveniles with IRF, NF-kB or STAT (red).
Second column (letter, one prime), CFDA-SE-labelled foreign or native bacteria (green).
Third column (letter, double prime), DNA in nuclei labelled with Hoechst (blue).
Fourth column (letter, triple prime), merged micrographs used in Fig. 4.
Scale bars, 10 µm.

**Fig. S10. Immunolocalization of IRF, STAT, and NF-κB in response to native and foreign bacteria at 1 hpe.**
(A, C, E) Separated fluorescence channels of panels shown in Fig. 4 of 1 hpe juveniles exposed to foreign bacteria.
(B, D, F) Separated fluorescence channels of panels shown in Fig. 4 of 1 hpe juveniles exposed to native bacteria.
First column (letter, no prime), immunostained juveniles with IRF, NF-kB or STAT (red).
Second column (letter, one prime), CFDA-SE-labelled foreign or native bacteria (green).
Third column (letter, double prime), DNA in nuclei labelled with Hoechst (blue).
Fourth column (letter, triple prime), merged micrographs used in Fig. 4.
Scale bars, 10 µm.

**Fig. S11. Immunolocalization of IRF, STAT, and NF-κB in *A. queenslandica* juveniles exposed to living and heat-killed native bacteria.**
(A, D, G) Separated fluorescence channels of panels show juveniles exposed to FSW at 2 hpe.
(B, E, H) Separated fluorescence channels of panels show juveniles exposed to heat-killed healthy native bacteria at 2 hpe.
(C, F, I) Separated fluorescence channels of panels show juveniles exposed to healthy native bacteria at 2 hpe.
First column (letter, no prime), immunostained juveniles with IRF, NF-kB or STAT (red).
Second column (letter, one prime), DNA in nuclei labelled with Hoechst (blue).
Third column (letter, double prime), merged micrographs.

(A-C) IRF is localised in the vicinity of choanocyte chambers (dotted lines) in juveniles treated with (A) filtered seawater, (B) heat-killed native bacteria and (C) living native bacteria. (C) IRF is present in numerous nuclei of amoebocytes (magenta arrowheads) and archaeocytes (large nuclei with a nucleoli; white arrowheads) in the mesohyl of juveniles exposed to living native bacteria.
(D-F) NF-κB is localised in the vicinity of choanocyte chambers (dotted lines) in juveniles treated with (D) filtered seawater, (E) heat-killed native bacteria and (F) living native bacteria. (E) NF-κB is weakly detected in some nuclei of amoebocytes and archaeocytes in heat-killed native bacteria, and (F) is present in numerous amoebocytes and archaeocytes in juveniles exposed to living native bacteria.
(G-I) STAT is localised in the vicinity of choanocyte chambers (dotted lines) in juveniles treated with (G) filtered seawater, (H) heat-killed native bacteria and (I) living native bacteria. (H) STAT is detected in some mesohyl cell nuclei of amoebocytes and archaeocytes in heat-killed native bacteria, and (I) is present in numerous amoebocytes and archaeocytes in juveniles exposed to living native bacteria.
Scale bars, 10 µm.

**Supplementary Tables**

**Table S1. Bacterial OTUs in healthy and unhealthy *Amphimedon queenslandica* and *Rhabdastrella globostellata*.** S1.1. OTU read counts obtained for bacterial communities enriched from *A. queenslandica* and *R. globostellata*. S1.2. Relative abundance of OTUs enriched from *A. queenslandica* and *R. globostellata*.
(XLSX)

**Table S2. CEL-Seq2 library statistics.** S2.1. Transcriptome mapping statistics for *A. queenslandica* juveniles exposed to filter seawater, native or foreign bacterial treatments. S2.2. Average transcriptome mapping statistics for *A. queenslandica* juveniles exposed to filter seawater, native or foreign bacterial treatments.
(XLSX)

**Table S3. Significantly differentially expressed protein coding genes identified by DESeq2.** Differentially expressed genes in juveniles exposed for 2 hours to foreign bacteria (S3.1), healthy native bacteria (S3.2) and unhealth native bacteria (S3.3). Differentially expressed genes in 8 hpe juveniles to foreign bacteria (S3.4), healthy native bacteria (S3.5) and unhealthy native bacteria (S3.6). Annotation of differentially expressed SCRCs (S3.7), GPCRs (S3.8) and IgSFs (S3.9); see Fig. S4 for relative expression levels. Protein coding sequences with no significant identity are not named.
(XLSX)

**Table S4. Relative expression levels of bacterial treatment-responsive genes in adult *A. queenslandica* cell types.** Adult sponges were collected from the wild and maintained in aquaria for less than week before cell dissociation [3, 4]. They were not treated with exogenous bacteria.

S5.1. Expression of 91 TFs in the 3 cell types that were manually curated and isolated. Collapsed vst counts are generated from 10 choanocyte, 15 archaeocyte and 6 pinacocyte transcriptomes [3]. S5.2. Expression of 85 TFs in the 13 cell types that were identified based on single-cell transcriptomes [4]. S5.3. Expression of 128 PRRs in the 3 cell types [3]. S5.4. Expression of 104 PRRs in the 13 cell types. S5.5. Expression of 53 immunity pathway genes in the 3 cell types. S5.6. Expression of 51 immunity pathway genes in the 13 cell types. S5.7. Expression of 41 xenobiotic metabolism genes in the 3 cell types. S5.8. Expression of 39 xenobiotic metabolism genes in the 13 cell types.
(XLSX)

**Table S5. Top protein coding and long non-coding RNA genes underlying differences between transcriptomes identified by sPLS-DA.** S4.1. 377 protein-coding genes and 3 lncRNAs contributing most to principal component 1. S4.2. 89 protein-coding genes and lncRNAs contributing most to principal component 2.
(XLSX)

**Table S6. Protein coding genes comprising six co-expression modules identified by WGCNA.** S6.1. Summary of 886 co-expressed genes in Module 1 (Blue). S6.2. Summary of 631 co-expressed genes in Module 2 (Brown). S6.3. Summary of 907 co-expressed genes in Module 3 (Turquoise). S6.4. Summary of 106 co-expressed genes in Module 4 (Pink). S6.5. Summary of 348 co-expressed genes in Module 5 (Yellow). S6.6. Summary of 113 co-expressed genes in Module 6 (Black).
(XLSX)

**Table S7. Intersection of WGCNA and sPLS-DA protein coding gene lists for modules 1-4.** S7.1. Summary of 268 principal component 1 feature genes in Module 1 (Blue). S7.2. Summary of 52 principal component 1 feature genes in Module 2 (Brown). S7.3. Summary of 27 principal component 2 feature genes in Module 3 (Turquoise). S7.4. Summary of 3 principal component 2 feature genes in Module 4 (Pink). Modules 1 and 2, and modules 3 and 4 are largely comprised of genes that are significantly differentially expressed in juveniles exposed to native and foreign bacteria, respectively.
(XLSX)
