## Supplementary figures and images for "Early transcription factor activation distinguishes symbiotic from non-symbiotic bacteria during microbiome processing in a sponge"

### Fig. S1.pdf

#### Supplementary Fig. S1

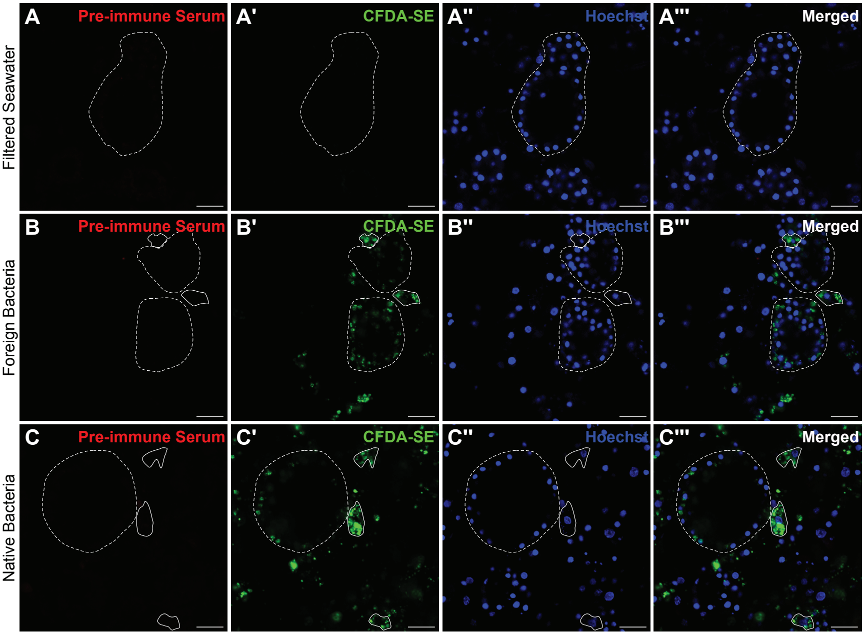

### Fig. S2.pdf

Supplementary Fig. S2

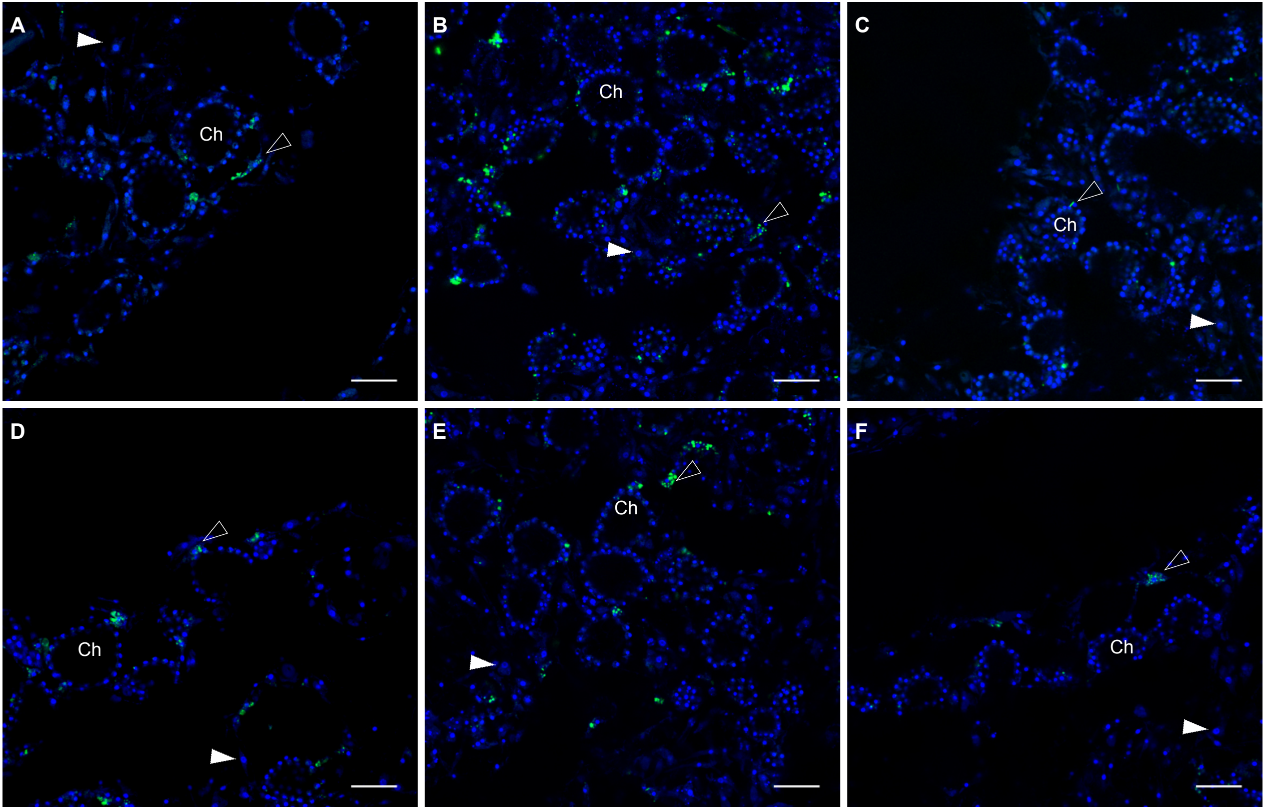

### Fig. S3.pdf

Supplementary Fig. S3

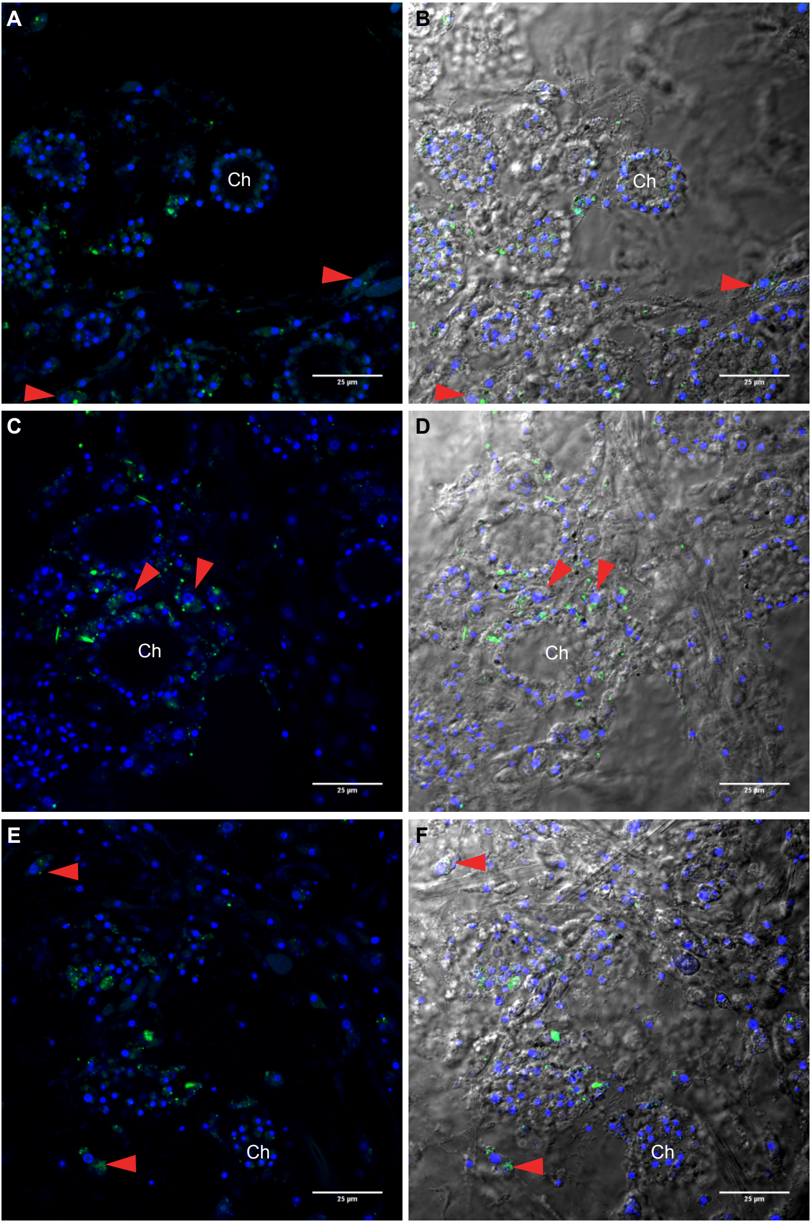

### Fig. S4.pdf

#### Supplementary Fig. S4

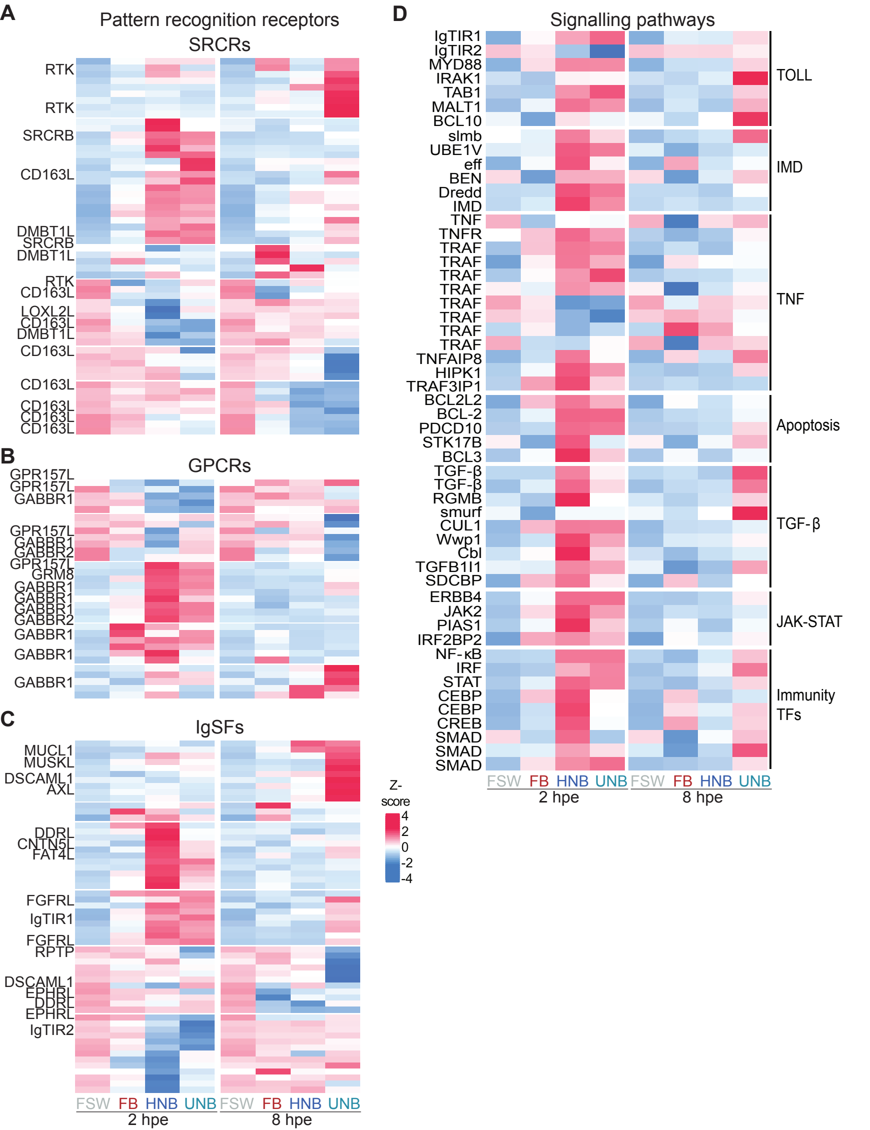

### Fig. S5.pdf

#### Supplementary Fig. S5

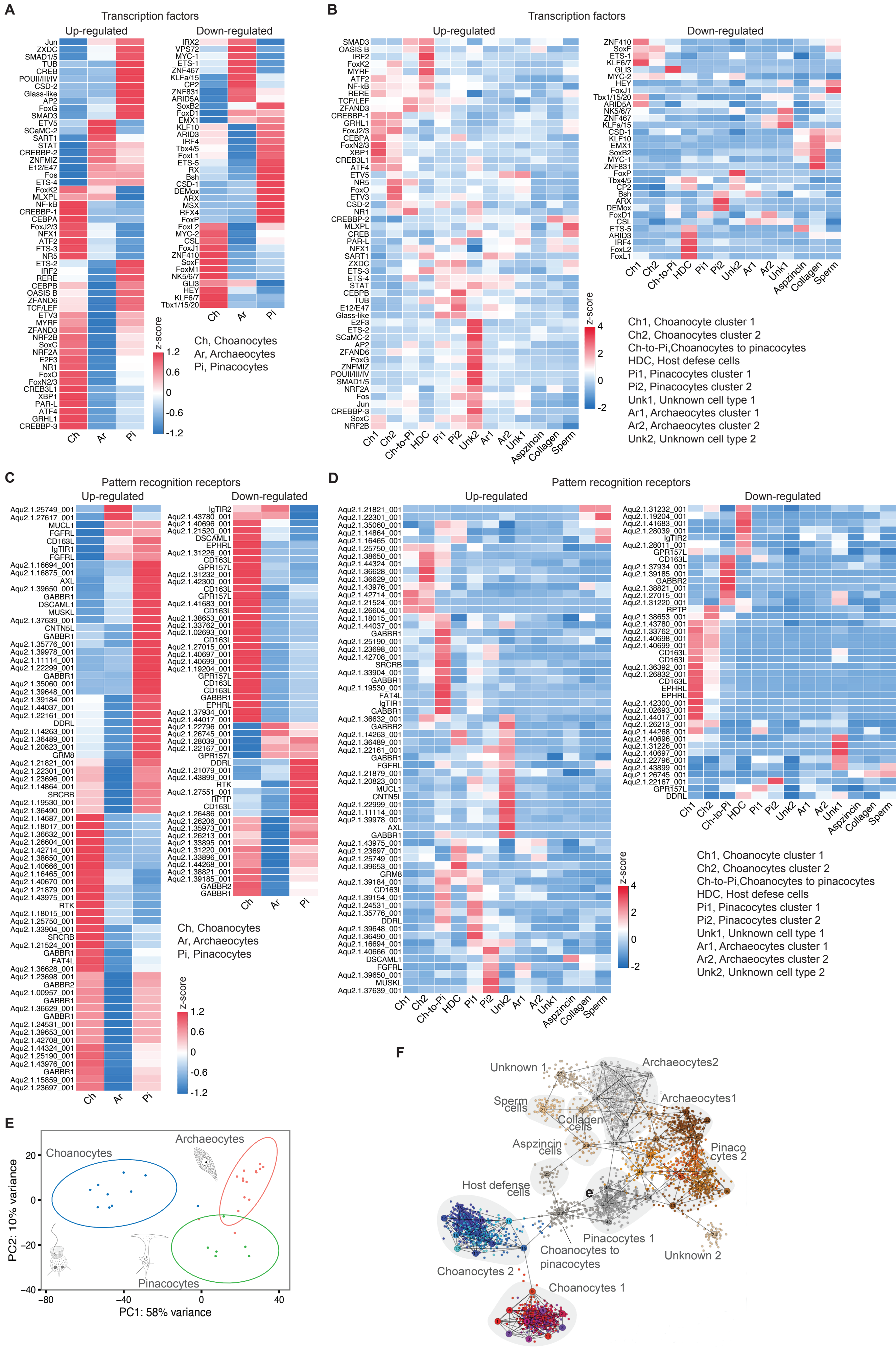

### Fig. S6.pdf

#### Supplementary Fig. S6

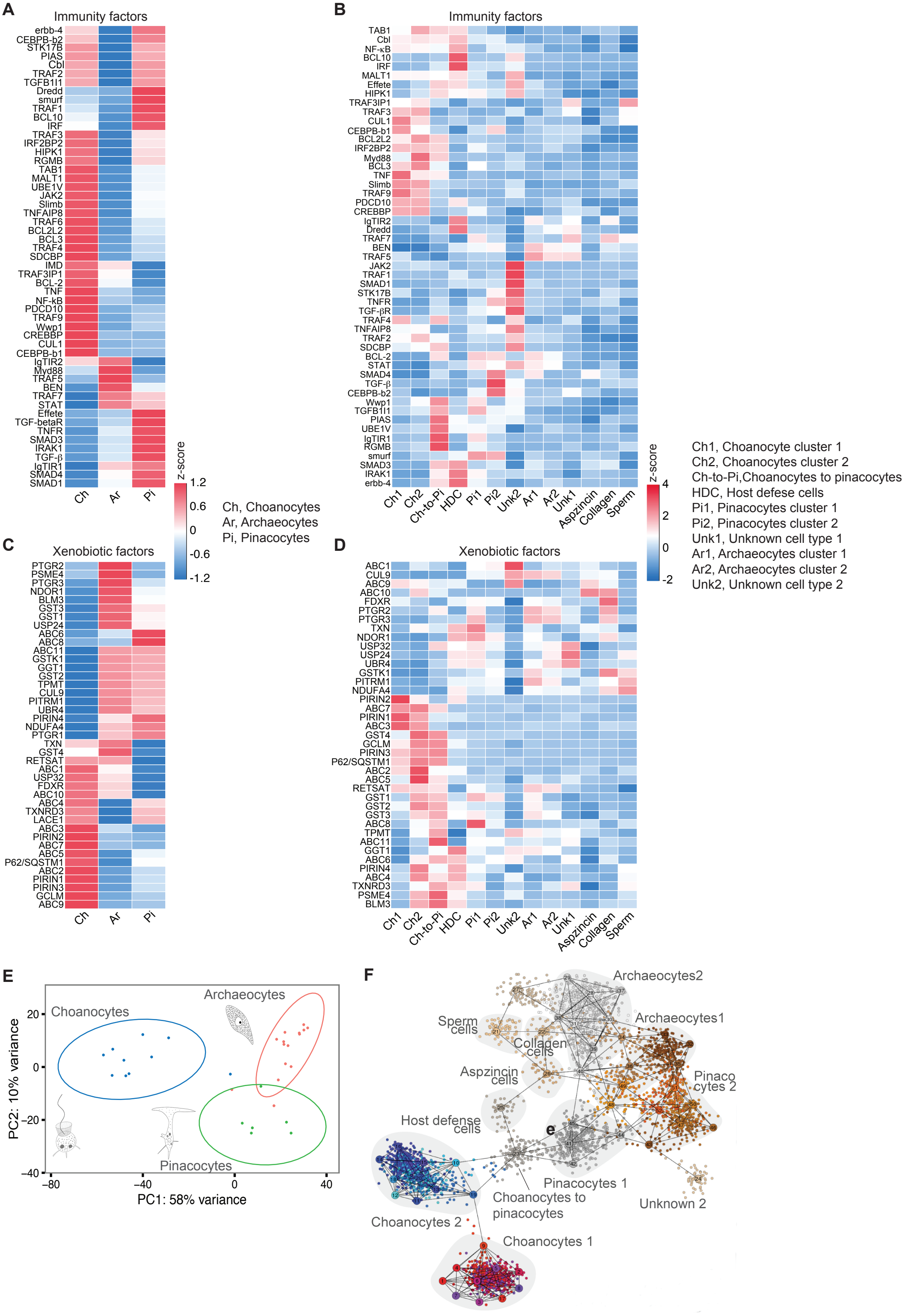

### Fig. S7.pdf

#### Supplementary Fig. S7

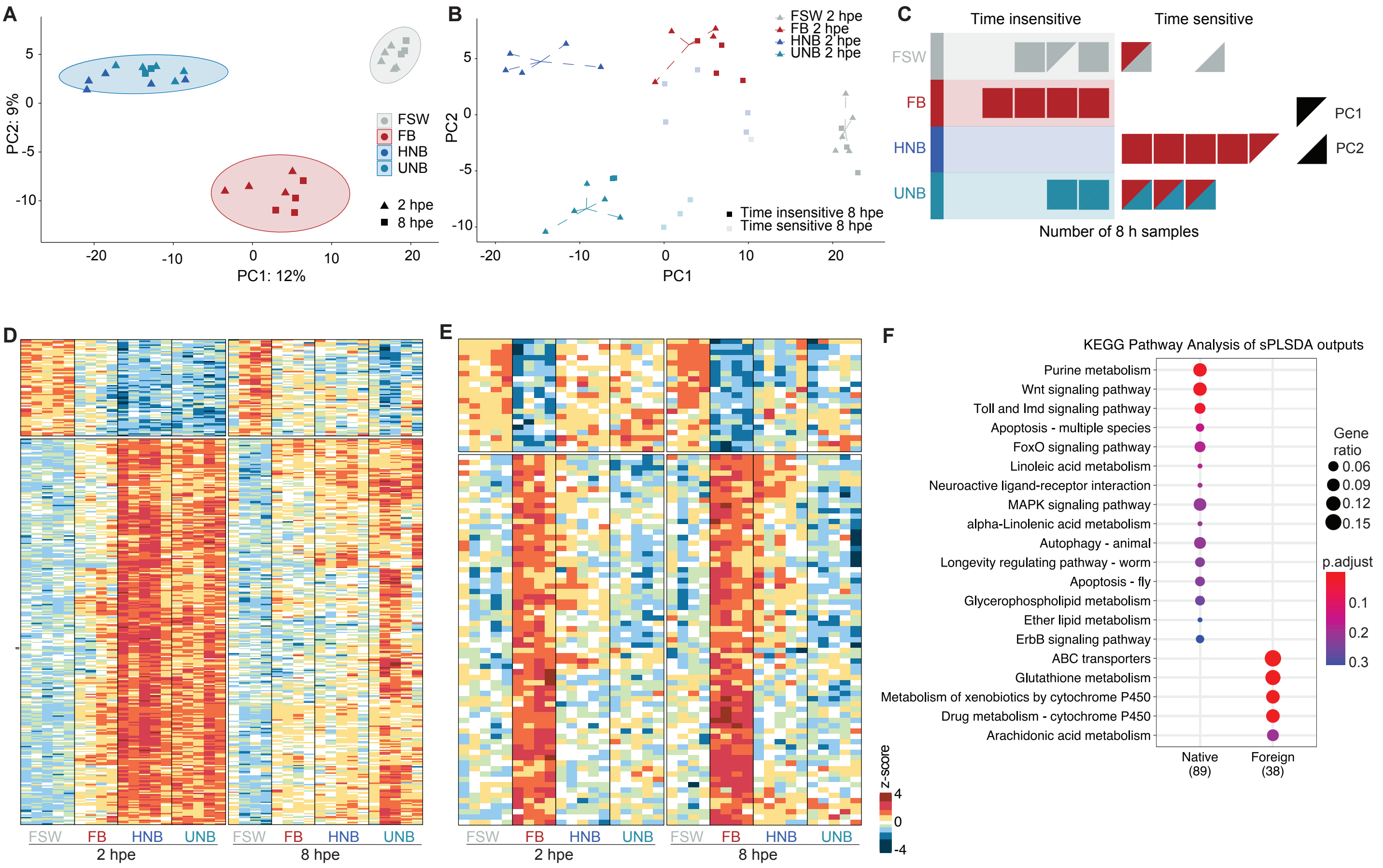

### Fig. S8.pdf

#### Supplementary Fig. S8

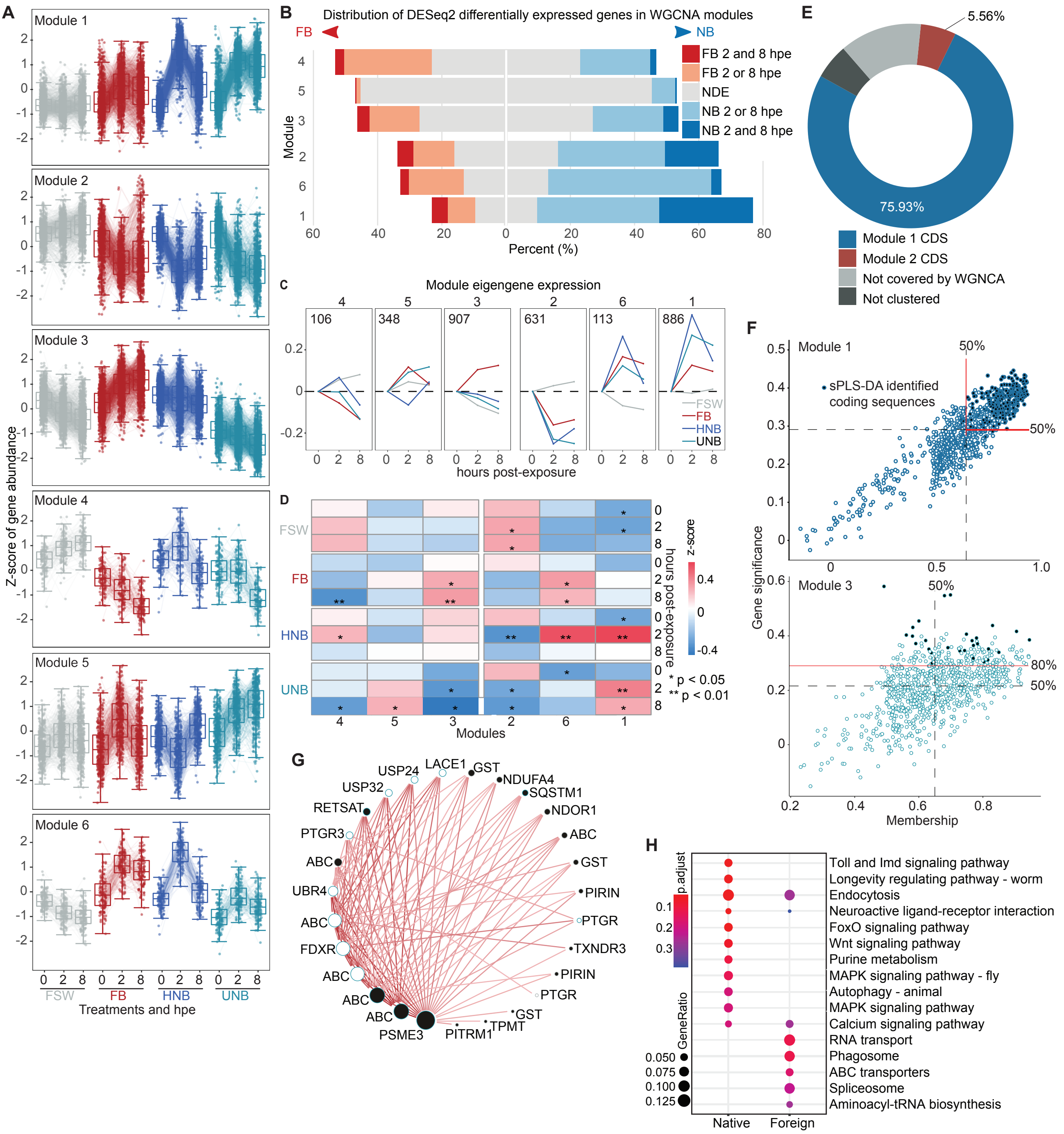

### Fig. S9.pdf

#### Supplementary Fig. S9

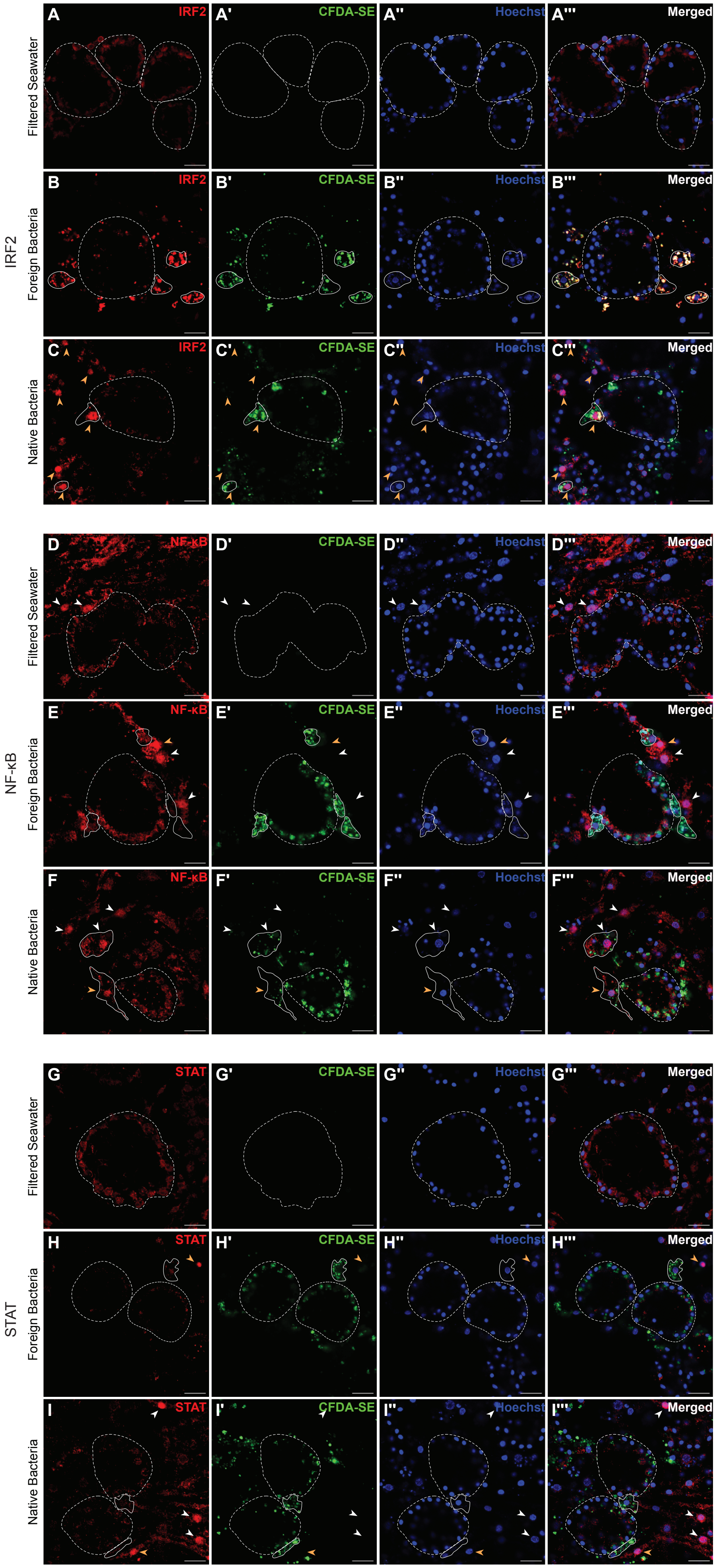

### Fig. S10.pdf

Supplementary Fig. S10

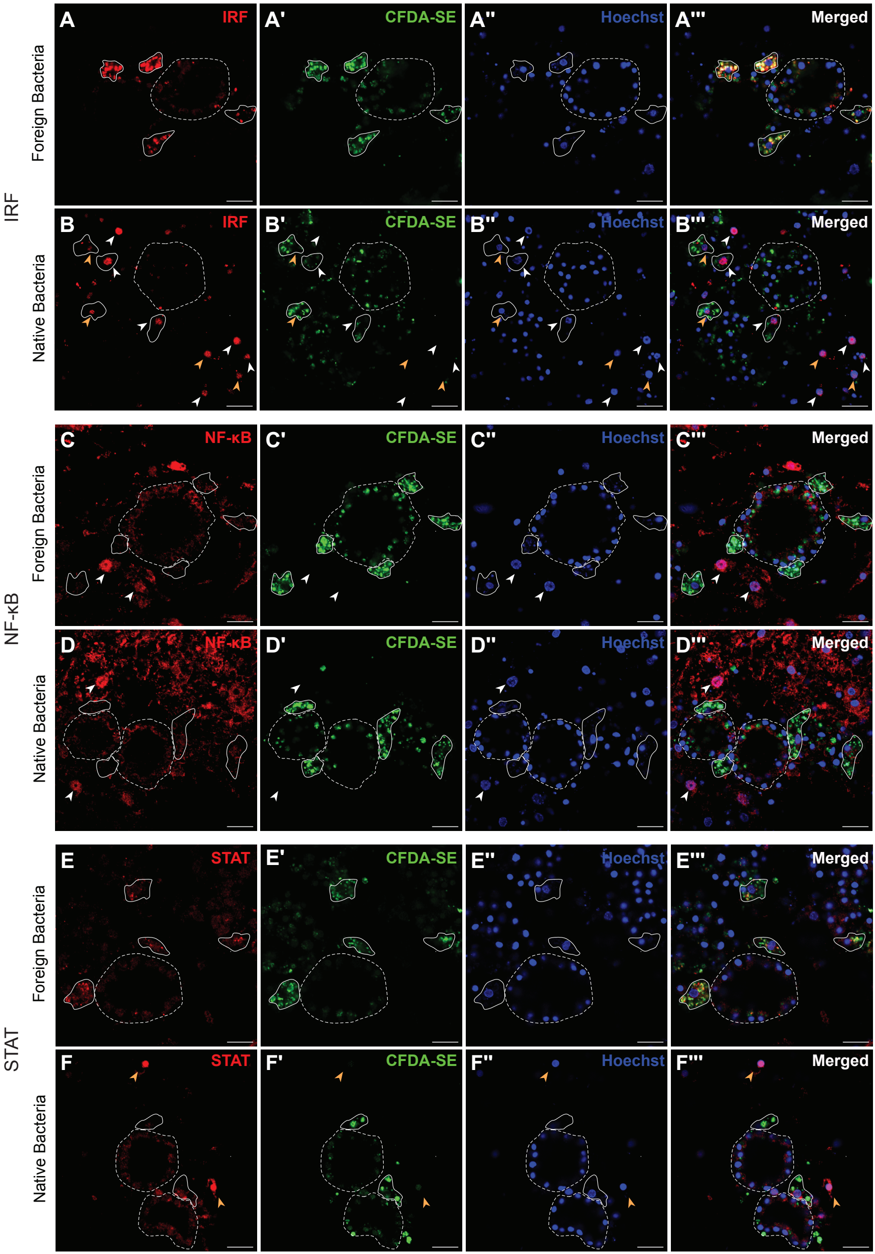

### Fig. S11.pdf

#### Supplementary Fig. S11

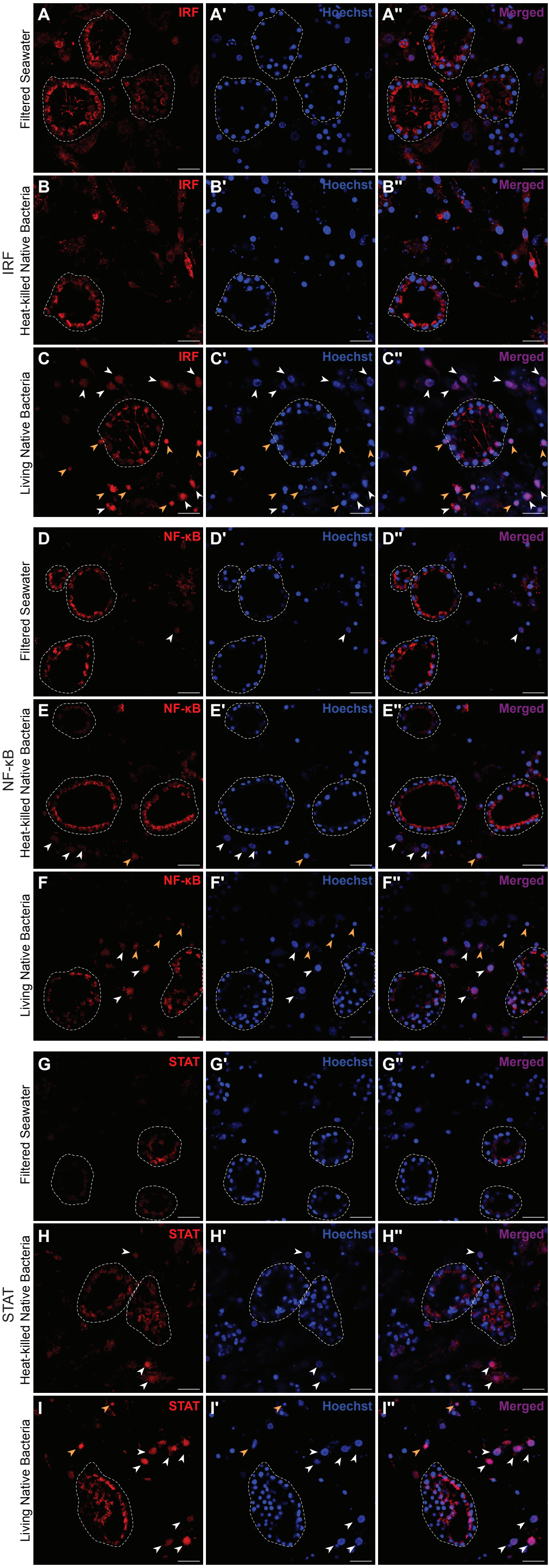
